# Nitrification in a Seagrass-Sponge Association

**DOI:** 10.1101/2025.01.07.631680

**Authors:** Johanna Berlinghof, Luis M. Montilla, Travis B. Meador, Luigi Gallucci, Donato Giovannelli, Harald Gruber-Vodicka, Maira Maselli, Francesca Margiotta, Christian Wild, Ulisse Cardini

## Abstract

In the Mediterranean Sea, the demosponge *Chondrilla nucula* can occur in close association with the native seagrass *Posidonia oceanica*. *C. nucula* harbors a diverse and abundant microbial community, including potential nitrifiers. Thus, the sponge may contribute to the nitrogen (N) demand of the seagrass holobiont. In this study, we investigated potential nitrification rates (PNR) and inorganic N fluxes within this association at a site where *C. nucula* covered 18 ± 3 % of the seagrass meadow area, during plant growth (spring) and senescence (autumn). Using incubation experiments with ^15^N-labeled ammonium, we measured PNR and inorganic N of the seagrass-sponge association, and of sponge and seagrass independently, under light and dark conditions. We supplemented these experiments with 16s rRNA gene amplicon sequencing to characterize the microbial community of the sponge. PNR was exclusively measured when the sponge was present (alone or in association with the seagrass). PNR was highest in the dark and when *C. nucula* was associated with the seagrass, ranging from 21 ± 7 to 267 ± 33 nmol N g DW^-1^ h^-1^ in spring and autumn, respectively. Sponge-mediated PNR can support 8% of the N demand of the *P. oceanica* holobiont during growth and 47 % during senescence. We identified key nitrifying bacterial and archaeal groups as members of the sponge’s microbial community. While *C. nucula* released inorganic N, potentially sustaining the seagrass, it benefitted from dissolved organic carbon released by *P. oceanica*. These results suggest that the interaction between *C. nucula* and *P. oceanica* is mutually beneficial, ultimately supporting and stabilizing the seagrass ecosystem.

## Introduction

Seagrasses are marine flowering plants that form vital ecosystems in coastal regions around the world, providing a range of ecological services and supporting high biodiversity (Costanza et al., 1997; Nordlund et al., 2016). Seagrasses can be considered as holobionts, forming complex symbiotic relationships with a diverse microbiome that includes bacteria, fungi, and other microorganisms living on the plant surfaces and within their tissues. These microbial communities are crucial for plant physiology and health because of their role in nutrient cycling, access to sunlight, or as protection against pathogens (e.g., Seymour et al., 2018; Tarquinio et al., 2019; Ugarelli et al., 2017). For instance, leaf epiphytes contribute to plant nitrogen (N) requirements by fixing atmospheric N_2_ and converting into bioavailable forms (e.g., Agawin et al., 2016; Mohr et al., 2021; Welsh, 2000), while sulfate-reducing bacteria in the rhizosphere contribute to nutrient mineralization (Holmer et al., 2001; Nielsen et al., 2001). The seagrass holobiont is further embedded in a ‘nested ecosystem’ (see Pita et al., 2018), where larger organisms, such as lucinid clams or sponges and their respective microbiome, interact with seagrasses and their associated microbes (Cardini et al., 2022; Malkin & Cardini, 2021). These nested interactions create a complex web of relationships that fundamentally contribute to the overall functioning of seagrass ecosystems.

Marine sponges (Porifera) represent one of the oldest and most primitive multicellular organisms on Earth. As filter feeders, they feed on microorganisms and can host dense and diverse microbial communities in their mesohyl matrix (Thomas et al., 2016). In recent years, there has been a growing emphasis on researching sponges and their prokaryotic symbionts, investigating their role in the biogeochemical cycling of nutrients (Maldonado et al., 2012 and references therein; Pita et al., 2018) and particularly on quantifying fluxes of dissolved inorganic nitrogen (DIN) released by sponges. Sponge-associated nitrification was hereby found to be the process producing the bulk of the DIN released (Diaz & Ward, 1997; Southwell, Weisz, et al., 2008).

Sponges are commonly found in seagrass meadows (Ávila et al., 2015; Soest et al., 2012) and can grow in very close association with the plants. However, the mechanisms and potential benefits of this association in terms of nutrient cycling still need to be investigated. Seagrasses are known to release large quantities of dissolved organic matter (DOM) into the surrounding seawater and sediments (Barrón & Duarte, 2009; Sogin et al., 2022). Sponges have the ability to take up DOM, which is then recycled to particulate organic matter (POM) that is released by the sponge and can be taken up by higher trophic levels. This process is also known as the “sponge loop”, a benthic counterpart of the oceanic microbial loop (De Goeij et al., 2013; Rix et al., 2017). The seagrass on the other hand may benefit from the release of DIN by the sponges via ammonium excretion, nitrogen fixation, or nitrification (Davy et al., 2002; Jiménez & Ribes, 2007; Fiore et al., 2010; Rix et al., 2015), as primary production in oligotrophic areas is often N-limited. Studies of an association between the seagrass *Thalassia testudinum* and the sponge *Halichondria melanadocia* in the Caribbean Sea revealed a context depended commensal relationship, balancing between the negative shading effect of the sponge for the seagrass with positive effects of N and phosphorus supplied by the sponge (Archer et al., 2015). This way, sponges can facilitate the growth of primary producers (Archer et al., 2021). At the same time, the sponge benefits from the substrate for growth provided by the plant (Archer et al., 2015).

A similar association can be found in the Mediterranean Sea between the demosponge *Chondrilla nucula* and the endemic seagrass *Posidonia oceanica*. The sponge can be found growing in very close association with the seagrass, attached to the lower part of the leaves. Belonging to the high microbial abundance (HMA) sponges, *C. nucula* harbors a distinct and diverse procaryotic community, including Cyanobacteria, Acidobacteria, Gamma-, and Deltaproteobacteria (Hill et al., 2006; Thiel et al., 2007) and also potential nitrifiers (Mazzella et al., 2024). Studies showed that *C. nucula* can release high amounts of DIN (17 - 44 nmol DIN g dry wt^-1^ min^-1^, Diaz & Ward, 1997; 141 ± 26 μmol NO_3_^-^ + NO_2_^-^ L^−1^ sponge h^−1^, Hoer et al., 2018; 600 nmol NO_3_^-^ dry wt^-1^ h^-1^, Corredor et al., 1988). The excretion of nitrate or nitrite is taken as first evidence of the presence of microbial nitrifiers in the sponges (Corredor et al., 1988; Diaz & Ward, 1997; Jiménez & Ribes, 2007). Understanding the mechanisms and rates of nitrification in these associations is important for unraveling the complexity of N dynamics in coastal ecosystems, with consequences for biodiversity and ecosystem functioning.

Nitrification is a pivotal process in the nitrogen (N) cycle and plays a fundamental role in shaping the nutrient dynamics of marine ecosystems. This biological transformation involves the oxidation of ammonia (NH_3_) to nitrite (NO_2-_) and subsequently to nitrate (NO_3_^-^), each process mediated by distinct groups of microorganisms (Ward, 2008). The first step, the oxidation from ammonia to nitrite, is performed by ammonia-oxidizing bacteria (AOB) or archaea (AOA), while the second step, the oxidation of nitrite to nitrate, is carried out by nitrite-oxidizing bacteria (NOB) (Ward, 2008). These nitrifying microorganisms can be found in the open ocean (Beman et al., 2013; Francis et al., 2005; Wuchter et al., 2006), coastal sediments (Freitag & Prosser, 2003; Park et al., 2008), but also marine invertebrates, such as sponges (Bayer et al., 2007; Hoffmann et al., 2009). Measurements of high nitrification rates based on the release of nitrite and nitrate have been reported from several tropical and temperate sponges (Bayer et al., 2008; Diaz & Ward, 1997; Hoffmann et al., 2009; Jiménez & Ribes, 2007; Nemoy et al., 2021).

In this study, we investigated the process of sponge-associated microbial nitrification as an indicator of a potential mutualism in the association between *P. oceanica* and *C. nucula*. We quantified potential nitrification rates (PNR) and net fluxes of inorganic and organic nutrients within the association and the organisms alone in incubation experiments, using ^15^N labeled ammonium, both in the light and in the dark. We complement these analyses with 16s rRNA gene amplicon sequencing to explore the diversity of the sponge microbial community, and the potential players involved in nitrification.

## Methods

### Study site and sampling

The incubation experiments were performed in May and October 2022 at the Schiacchetiello inlet (40°47’36.9″N 14°05’13.4″E) in the area of Bacoli (Tyrrhenian Sea, Italy). Here, shallow patches (0-6 m depth) of a *P. oceanica* meadow with a high *C. nucula* coverage exist (Table 1). The site is characterized by high human pressure due to tourism (e.g., boat anchoring in the meadows) and eutrophication due to a nearby mussel farm. We collected the aboveground part of *P. oceanica* shoots when growing alone, small specimens of *C. nucula* (max. 5 cm ∅) growing alone and both when growing in association. We selected *P. oceanica* shoots in the central part of the meadow patches to avoid edge effects. *C. nucula* was carefully removed from the substrate to avoid any damage to the tissue. Shoots of *P. oceanica* with the sponge growing attached to the lower part of the leaves were considered as association. We made sure that the organisms stayed submerged in the water until further use.

**Table 1.** Environmental parameters (mean ± SE) measured in May and October 2022. Temperature and light were continuously measured with data loggers (between ca. 10 am and 5 pm of the respective incubation day). DOC, DON, NH_4_^+^, and NO_3_^-^ were analyzed from samples collected on the respective sampling day (n = 3). Sponge coverage and shoot density were collected in autumn at 8 meadow patches with 8 random subplots each (25 x 25 cm).

|  | Spring | Autumn |
| --- | --- | --- |
| Water temperature ($^{\circ}\text{C}$) | $21.03 \pm 0.01$ | $24.52 \pm 0.04$ |
| Light (Lux) | $58441 \pm 2748$ | $39007 \pm 1498$ |
| Dissolved oxygen (mg/L) | $8.10 \pm 0.03$ | $7.52 \pm 0.12$ |
| DOC ($\mu\text{M}$) | $106.46 \pm 4.91$ | $134.37 \pm 8.03$ |
| DON ($\mu\text{M}$) | $7.82 \pm 0.68$ | $11.25 \pm 0.29$ |
| $\text{NH}_4^+$ ($\mu\text{M}$) | $12.74 \pm 3.09$ | $2.76 \pm 1.75$ |
| $\text{NO}_3^-$ ($\mu\text{M}$) | $2.06 \pm 1.23$ | $2.21 \pm 0.03$ |
| Sponge coverage (%) | --- | $18.33 \pm 2.72$ |
| Seagrass shoot density ($\text{m}^{-2}$) | --- | $349.00 \pm 16.71$ |

Samples for the microbial community analysis of the sponge were collected in May 2022 at the site described above. Five specimens of *C. nucula* growing alone and in association were removed from the substrate, cut into pieces with sterile scalpels, and washed with sterile filtered seawater to remove rubble or organisms attached. They were then transferred into sterile 50 mL Falcon tubes filled with stabilizing buffer solution (RNA*later*) and stored on dry ice until transferred to the laboratory, where they were stored at −20°C until further analysis. For the microbial community of the water column, we collected 5 L of seawater from the sampling site and filtered 5 x 1 L on 0.22 μm cellulose nitrate membrane filters. The filters were transferred into sterile 15 mL Falcon tubes filled with RNA*later* and stored on dry ice until transfer to the laboratory, where they were stored at −20°C until further analysis.

### Incubation experiment with stable isotopes

PNR was determined by amending site water with 5 μM ^15^NH_4_^+^ (≥98 atom %^15^N) tracing solution. For the incubations, we filled acid-washed polyethylene chambers (1100 mL) with site water and added seagrass, sponge, or the association (n = 4) for incubating in the light or the dark. We added 500 µL of a 10 mM ^15^NH_4_^+^ stock solution to each chamber, closed them without air bubbles, and gently inverted them. This resulted in an enrichment of 60.19 atom% ^15^N-NH_4_^+^ in spring and of 76.51 atom% in autumn in the incubation chambers. ^15^N-NH_4_^+^-enriched chambers (also 5 μM) without organisms served as controls for background processes in the water column (n = 2). Chambers with the seagrass-sponge association but without ^15^N-NH_4_^+^- enrichment served as controls for our isotope enrichment method (n = 2).

T_0_ samples for the analysis of O_2_ production were taken from the bottom of the chambers to reduce gas exchange in 12 mL exetainers (Labco Ltd.) using acid-washed syringes and tubes and were subsequently fixed with 100 μL 7 M ZnCl_2_. T_0_ samples for the analysis of dissolved inorganic nutrients (DIN: NH_4_^+^, NO_2_^-^, NO_x_^-^ and PO_4_^3-^), dissolved organic carbon and nitrogen (DOC, DON), and ^15^NO_3_^-^ concentrations were taken in triplicates from an extra incubation chamber with 5 μM ^15^N-NH_4_^+^-enrichment without organisms added. The chambers were refilled with site water, closed without air bubbles, and placed upside down in two plastic crates, making sure the seagrass shoots and sponges were placed the right way around and to not create shading from the chamber lids or the crates. One crate was covered with black plastic bags for the dark incubation; then both crates were placed for 5-6 h floating in the water to ensure a stable temperature, light availability, and mixing via wave activity (16 chambers per crate, 32 chambers in total). Temperature and light were continually measured during the incubation with HOBO data loggers (Onset Computer Corporation) inside a control chamber and in the water column.

At the end of the incubation, the chambers were opened and T_final_ samples for O_2_ analysis were taken as described above. Samples for the analysis of ^15^NO_3-_ production were filtered with 0.2 μm disposable syringe filters into 50 mL acid-washed Falcon tubes and stored on dry ice until transported to the laboratory, where they were frozen at −20°C until further analysis. Samples to analyze DIN were filtered with 0.2 μm disposable syringe filters into 20 mL acid-washed HDPE vials and stored on dry ice until transported to the laboratory, where they were frozen at −20°C until further analysis. Samples to analyze DOC and DON were filtered with precombusted (400°C, 4 to 5 h) 0.7 μm GF/F filters into 30 mL acid-washed HDPE vials, fixed with 80 μL 18% HCl, and stored in cool boxes until transferred to the laboratory, where they were stored at 6°C until further analysis. The biomass (seagrass and sponges) was collected in zip-lock bags and stored in cool boxes until further processing. In the lab, the biomass samples were then processed by separating seagrass leaves, seagrass epiphytes, and sponges for measuring the wet weight. After the samples were freeze-dried for 24 h, the dry weight of each tissue type was measured. To measure background C and N concentrations at the study site, 1000 mL of water column samples for T_0_ as well as the remaining water of the incubation chambers at T_final_ was filtered with precombusted 0.7 μm GF/F filters. The filters were stored in 15 mL centrifuge tubes on dry ice until transport to the laboratory, where they were frozen at −20°C until further analysis.

### Potential nitrification rates (PNR)

Isotopic samples for ^15^NO_3_^-^ production were analyzed by isotope ratio mass spectrometry (IRMS, ThermoScientific) using the Ti(III) reduction method as described in (Berlinghof et al., 2024). Potential nitrification rates (PNR) were calculated using an equation modified from (Beman et al., 2011):

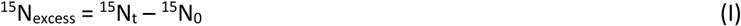

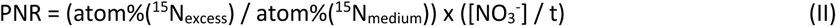

^15^N_t_ is the ^15^N content of the samples in the NO_3_^−^ pool measured at time t, and ^15^N_0_ is the ^15^N content in the NO_3_^−^ pool measured at the beginning of the incubations. The enrichment of samples (^15^N_excess_) was considered significant for samples with a value greater than 2.5 times the standard deviation of the mean of the T_0_ samples. ^15^N_medium_ is the enrichment of the incubation medium at the end of the incubations. Based on the NH_4_^+^ concentrations measured before and after the addition of ^15^NH_4_^+^, this resulted in an enrichment of ∼95.9 atom %^15^N in the incubation medium. [NO_3_^-^] is the concentration of NO_3_^-^ (μM) and t is the incubation time (h). PNR was corrected for the rates in control incubations without organisms. Since PNR was only detected in incubations where the sponge was present, they were normalized to sponge dry weight (g) in the SG + SP and SP treatments; and to seagrass dry weight (g) in the SG treatments.

### Stable Isotope Analysis

We collected samples of *P. oceanica* (leaves and epiphytes, in autumn additionally seagrass meristem tissue) and *C. nucula*, when growing associated vs non-associated. Epiphytes were carefully scraped off the seagrass leaves and stored in Eppendorf tubes. All samples were stored in acid-washed vials and lyophilized. The samples were ground to a fine powder using a tissue lyser, acid-fumed with HCl and then weighed into silver capsules for isotope analysis. The samples were analyzed using a Flash Elemental Analyzer (Thermo Scientific) equipped with a single reactor (1020°C), along with a MAT 253 Plus isotope ratio mass spectrometer (IRMS) interfaced with a Conflo IV system (Thermo Scientific, Bremen, Germany). The δ¹³C and δ¹⁵N values were corrected for blanks, ion source linearity, standardized against laboratory working standards and international reference materials (IAEA-600, IAEA-603) and normalized to Vienna Pee Dee Belemnite and atmospheric air, respectively. Precision was typically <0.1‰ for δ¹³C and 0.5‰ for δ¹⁵N. The molar C:N ratios (mol:mol) were calculated from C and N weights in the capsules (µg) and based on their respective molecular weights.

### Fluxes of oxygen, inorganic and organic nutrients

Oxygen concentrations from T_0_ and T_final_ sampling were measured with a membrane-introduction mass spectrometry (MIMS, Bay Instruments, LLC). All samples were measured in technical quadruplicates and 0.2 μM filtered seawater (20°C, salinity = 38 psu) was used as standard to calculate the O_2_ concentrations from the atomic mass of 32. The lowest oxygen saturation in the dark incubations dropped to 38% of the initial O_2_ concentration and the highest in the light incubation increased to 179%.

Dissolved inorganic nutrient concentrations (NH_4_^+^, NO_2_^-^, NO_3_^-^, PO_4_^-^) were measured with a continuous flow analyzer (Flowsys, SYSTEA SpA.). NO_3_^-^ concentrations were calculated as the difference between NO_x_^-^ and NO_2_^-^. DOC and DON concentrations were measured with a TOC-L Analyzer with TN unit (Shimadzu Corporation, Japan). Net nutrient fluxes were calculated as the difference between final and initial nutrient concentrations, corrected for controls, and normalized to biomass dry weight.

### Data analysis

To compute daily, integrated rates of nutrient fluxes (daily flux = light flux × 12 + dark flux × 12), we generated analytical combinations of the observed light and dark fluxes, assuming a daily 12:12 h light/dark cycle. Each pair of independent values was combined to calculate the distribution of integrated rates (n=4). The results present a thorough distribution of potential outcomes derived from the input data. Permutation-based analysis of variance (PERMANOVA) using Euclidean distance was performed on each response variable (Anderson, 2017) to test the effects of *community* (seagrass, sponge, association) and *season* (spring vs autumn) on PNR, O_2_, inorganic and organic nutrient fluxes. δ¹³C, δ¹⁵N values and C:N ratios were tested for differences among *tissue* types (*P. oceanica* leaves, *P. oceanica* meristem, *P. oceanica* epiphytes, *C. nucula*), *association* types (associated vs non-associated) and *season* (spring vs autumn) using PERMANOVA (n=8). Pairwise comparisons were performed using Tukey’s honest significant difference (HSD) test. All statistical analyses were performed with R version 4.2.3 (R Core team, 2023) using the packages *car* and *emmeans*.

### Holobiont N demand calculations

To calculate how much DIN *C. nucula* can provide via nitrification and ammonification for the N demand of the *P. oceanica* holobiont (plant + epiphytes), we integrated PNR in light and dark incubations assuming a daily 12:12 h light/dark cycle. We further used the daily O_2_ budget (using a photosynthetic quotient of 1) and C:N ratios of seagrass leaf and epiphyte tissue to calculate the potential percentage of daily primary production of the seagrass holobiont that can be supported by sponge-mediated PNR.

### Prokaryotic DNA extraction, amplification, and sequencing

DNA from sponge and seawater samples was extracted using the Qiagen DNeasy Powersoil Kit (Qiagen) following a modified version of the method described by Taylor et al. (2004). Sponge tissue was grounded, resuspended in sterile distilled water and left for 1 hour. The tissue was transferred into a fast-prep tube (or tube containing 0,5 g of silica beads) and 1 ml of extraction buffer, 0.015 g of PVPP, 300 microliters of chloroform-Isoamylic (24:1) was added. The fast-prep tube was centrifuged at 15.000 g for 30 min, the supernatant was collected and precipitated over night at room temperature with 3M sodium acetate (0.1 x sample volume) and isopropanol (0.7 x sample volume). Then the samples were centrifuged at 14.000 rpm for 30 min, and the pellets were washed twice with 70% ethanol, dried at 37°C, and re-suspended in 50 µl Tris HCL (pH 8; 10 mM). The water filters were cut in little pieces and transferred sterile into a 50 ml Falcon tube. Then 2.25 ml of extraction buffer and 100 microliter of Proteinase K (Stock 100 µg/µl) was added, and the samples were incubated at 37°C on a shaker for 30 minutes and then at 55°C for 30 min in a water bath. 0.25 ml of SDS 20% were added to each sample and they were incubated for 5 minutes in dry ice or −80° and 3x5 minutes in a water bath at 65°C. The samples were centrifuged for 10 minutes at 7000 rpm, and the supernatant was transferred into a new sterile 50 ml Falcon tube. 900 µl extraction buffer and 100 µl of SDS 20% were added to the pellet in the old Falcon tube, vortexted, incubated for 5 minutes at 65°C, centrifuge for 10 minutes at 7000 rpm, and the supernatant was transferred to the supernatant previously taken. Chloroform:Isoamyl alcohol (24:1; 1 x sample volume) was added, centrifuged for 10 minutes at 7000 rpm, and the supernatant was transferred into a new sterile 15 ml Falcon tube. Isopropanol (0.6 x sample volume) was added and precipitate overnight. Then each sample was splitted into 3 x 2 mL Eppendorf tubes, centrifuged for 30 minutes at 14000 rpm, the supernatant was discarded, the pellet was dried at room temperature for 1 hour and then resuspended in 50 µl of sterile water.

The extracted DNA samples were quantified using a microvolume spectrophotometer (Thermo Scientific NanoDrop 2000c) and stored at −20 °C until processing. PacBio Sequel sequencing of the full 16S rRNA gene was performed using the 27F (=AGRGTTYGATYMTGGCTCAG) and 1492R (=RGYTACCTTGTTACGACTT) bacteria-specific primers. Additionally, PacBio Sequel sequencing was performed with Arch21Ftrim (=TCCGGTTGATCCYGCCGG) and A1401R (=CRGTGWGTRCAAGGRGCA) as archaea-specific primers. The primers were removed from the raw sequence data and the fastq files were processed using the R package DADA2 v.1.28.0 (Callahan et al., 2016). Quality filtering and denoising of the trimmed fastq files was performed using the following parameters: “minQ=3, minLen=1000, maxLen=1600, maxN=0, rm.phix=FALSE, maxEE=2). Paired-end reads were then merged into amplicon sequence variants (ASVs); chimeric sequences were identified and removed. Prokaryotic taxonomy assignment was performed using the SILVA v 138.1 database.

### Bioinformatics and data analysis of the sequencing data

The ASV matrix was analyzed using the R package phyloseq v.1.44.0 (McMurdie & Holmes, 2013). Chloroplast and mitochondrial sequences were removed, the data was transformed to relative abundances and samples were pooled per treatment (sponge alone, sponge from association, water column) to calculate the average relative abundances. We tested the effects of sample type (sponge vs water column) and sponge type (associated vs non-associated) on the microbial community associated with *C. nucula* at genus level in a differential abundance analysis using the R package DESeq2 v.1.40.2 (Love et al., 2014). The dataset was then filtered for nitrifying taxa, and we tested the effects of sample and sponge type using a permutation-based multivariate analysis of variance derived from a Euclidean distance matrix using the vegan package v. 2.6.4 (Oksanen et al., 2020).

## Results

### Potential nitrification rates (PNR)

We explored the nitrification potential of the seagrass and sponge microbiomes in incubation experiments with amended ^15^N-NH_4_^+^. We found significant (>2.5 x SD of T_0_) potential nitrification rates (PNR) in incubations where the sponge was present but not when only seagrass was present (Fig. 1, Table S1). PNR were highest in the association, followed by the sponge, although differences between these treatments were only significant in autumn (Fig. 1, Table S2). In spring, PNR reached 5.31 ± 0.75 nmol g DW^-1^ h^-1^ (mean ± SE) in the association in the light (Fig. 1a) and 21.51 ± 6.76 nmol g DW^-1^ h^-1^ in the dark (Fig. 1b). In autumn, PNR in the association reached 106.15 ± 16.21 nmol g DW^-1^ h^-1^ in the light (Fig. 1c), and 267.25 ± 33.01 nmol g DW^-1^ h^-1^ in dark incubations treatment (Fig. 1d). The sponge showed intermediate PNR rates in spring (3.74 ± 1.93 nmol g DW^-1^ h^-1^ in the light and 13.40 ± 3.53 nmol g DW^-1^ h^-1^ in the dark, Fig. 1a, b) and in autumn (26.13 ± 6.85 nmol g DW^-1^ h^-1^ in the light and 45.39 ± 14.86 nmol g DW^-1^ h^-1^ in the dark, Fig. 1c, d). The seagrass showed PNR close to zero in all incubations (Fig. 1). PNR of the sponge and the association was higher in dark incubations than in the light with rates 259 % (sponge) and 305 % (association) higher in spring and 74 % (sponge) and 152 % (association) higher in autumn (Fig. 1, Table S2). A seasonal effect was particularly evident in the association, where PNR in autumn was one magnitude higher than in spring (Fig. 1, Table S2).

**Fig. 1.**
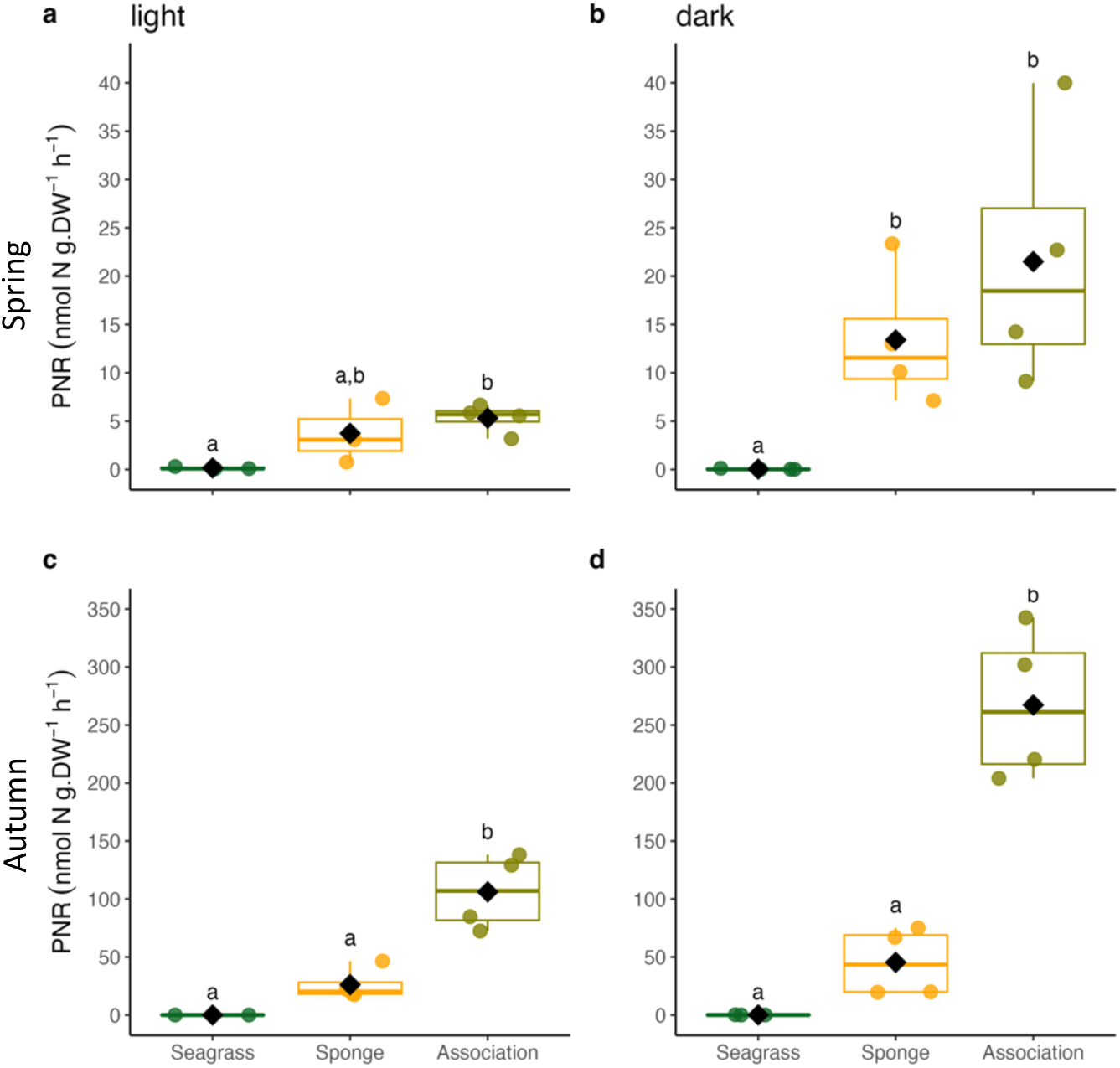
PNR of the seagrass, sponge and the association during light (a, c) and dark (b, d) incubations in spring and autumn. The center line denotes the median value (50^th^ percentile), and the box contains the 25^th^ to 75^th^ percentiles. Whiskers mark the 5^th^ and 95^th^ percentiles. Black squares indicate mean values; letters indicate significant differences between treatments, n=4.

### Stable Isotope Analysis

The stable isotope analysis of natural abundance samples in spring, revealed an increase in δ^15^N in old *P. oceanica* leaves and epiphytes living associated with the sponge, from 3.8 ± 0.4 ‰ to 5.6 ± 0.2 ‰ and from 5.9 ± 0.3 ‰ to 7.3 ± 0.5 ‰, respectively (Fig. S1a, Tables S4, S5). Conversely, the association type had no effect in autumn, but δ^15^N values were generally lower than in spring (Fig. S1b, Tables S4, S6). δ^13^C of the seagrass leaves ranged from −14.3 ± 0.2 ‰ in spring and −13.7 ± 0.2‰ in autumn, with no statistical differences between associated and non-associated state (Fig. S1c, d, Tables S4, S7, S8). δ^13^C values of the sponge *C. nucula* were higher in autumn than in spring (−19.1 ± 0.1 ‰ vs 18.4 ± 0.1 ‰, Fig. S1c, d) but showed no differences between associated and non-associated state (Tables S4, S7, S8). Conversely, δ^15^N values *C. nucula* were lower in autumn than in spring (6.6 ± 0.2 ‰ vs 5.1 ± 0.1 ‰, S1a, b), but also showing no effects of the association type (Tables S4, S5, S6). Plant epiphytes in spring had δ^13^C values similar to those of the sponge (−18.9 ± 0.4 ‰, Fig. S1c, d) and in autumn similar to seagrass leaves (−14.3 ± 0.5 ‰, Fig. S1c, d, Tables S4, S7, S5.8). Epiphytes showed an increase in their δ^15^N of 1.4 ‰ when the plant was associated with the sponge in spring (from 5.9 ± 0.3 ‰ to 7.3 ± 0.5 ‰, Fig. S1a, Table S4, S5), but not in autumn (Fig. S1b, Table S4, S6).

### Inorganic Nutrient Fluxes

Daily net NH_4_^+^ fluxes in spring showed a trend towards release by the sponge and uptake by the seagrass (14.71 ± 8.38 and −11.75 ± 4.88 μmol g DW^-1^ d^-1^, respectively, mean ± SE), while the association showed net fluxes close to zero (Fig. 2a, Tables S9, S10). Conversely, in autumn, all community types showed similar uptake rates (−19.68 ± 2.88 μmol g DW^-1^ d^-1^, Fig. 2b, Tables S9, S10). We observed daily net NO_3_^-^ production by sponges in spring and autumn (17.78 ± 3.49 and 23.59 ± 2.89 μmol g DW^-1^ d^-1^, respectively, Fig. 2c, d, Tables S9, S10). In spring, the seagrass and the association showed net fluxes close to zero (Fig. 2c, d). Conversely, in autumn, the seagrass showed NO_3_^-^ uptake, while we observed intermediate net production in the association (−11.23 ± 3.03 and 9.73 ± 3.03 μmol g DW^-1^ d^-1^, respectively, Fig. 2d, Tables S9, S10). We observed daily net NO_2_^-^ release in all incubations in spring, highest in the sponge (0.60 ± 0.02 μmol g DW^-1^ d^-1^), followed by the seagrass (0.28 ± 0.11 μmol g DW^- 1^ d^-1^) and the association (0.19 ± 0.12 μmol g DW^-1^ d^-1^, Fig. 2e, Tables S9, S10). In autumn, the seagrass and the association showed net NO_2_^-^ fluxes close to zero and the sponge a trend towards NO_2_^-^ release (0.15 ± 0.06 μmol g DW^-1^ d^-1^), but differences were not significant (Fig. 2f, Tables S9, S10).

**Fig. 2.**
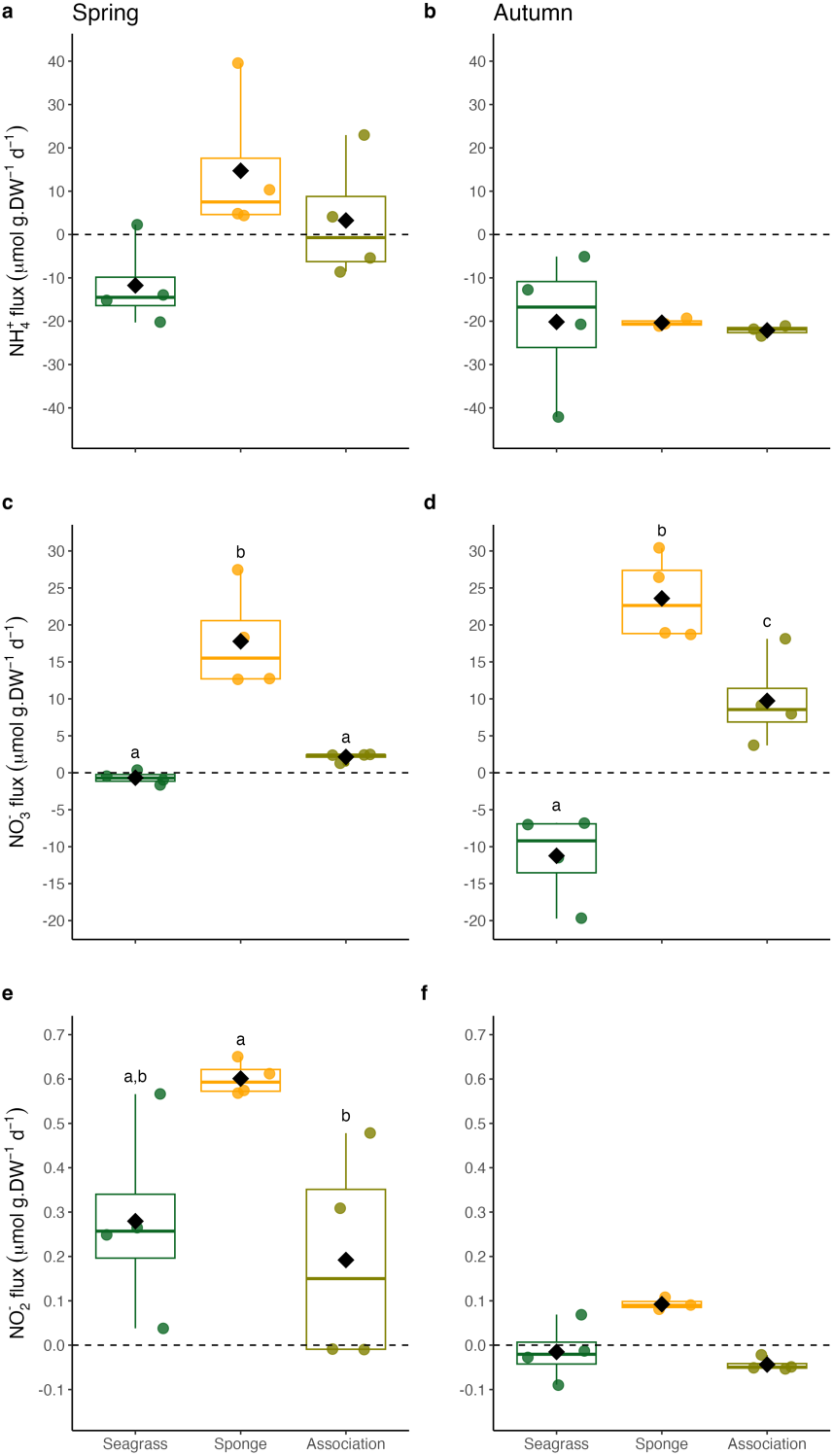
Daily fluxes of NH_4_^+^ (a, b), NO_3_^-^ (c, d), and NO_2_^-^ (e, f) in incubations with the seagrass, sponge and the association in spring and autumn. Positive values indicate production, negative values indicate uptake. The center line denotes the median value (50^th^ percentile), and the box contains the 25^th^ to 75^th^ percentiles. Whiskers mark the 5^th^ and 95^th^ percentiles. The horizontal dashed line marks the zero-flux threshold. Black squares indicate mean values; letters indicate significant differences between treatments, n=4.

### Organic Nutrient Fluxes

The seagrass was a constant source of daily net DOC flux in spring and in autumn (140.05 ± 32.35 and 185.10 ± 36.97 μmol g DW^-1^ d^-1^, respectively; mean ± SE, Fig. 3a, b, Tables S11, S12). We observed net DOC fluxes close to zero of the sponge in spring and in autumn, while the association showed net fluxes close to zero in spring and uptake in autumn (−40.37 ± 15.35 μmol g DW^-1^ d^-1^, Fig. 3a, b). Conversely, the sponge was a constant source of daily net DON flux in spring and in autumn (38.90 ± 5.36 and 54.42 ± 2.84 μmol g DW^-1^ d^-1^, respectively, Fig. 3c, d, Tables S11, S12). The seagrass and the association showed similar net DON production in spring (5.84 ± 3.80 and 9.07 ± 5.05 μmol g DW^-1^ d^-1^, respectively, Fig. 3c). In autumn, the seagrass showed DON production (16.16 ± 5.72 μmol g DW^-1^ d^-1^, Fig. 3d) and the association net fluxes close to zero (Fig. 3d). Seasons had no significant effect on daily net DOC or DON fluxes (Tables S11, S12).

**Fig. 3.**
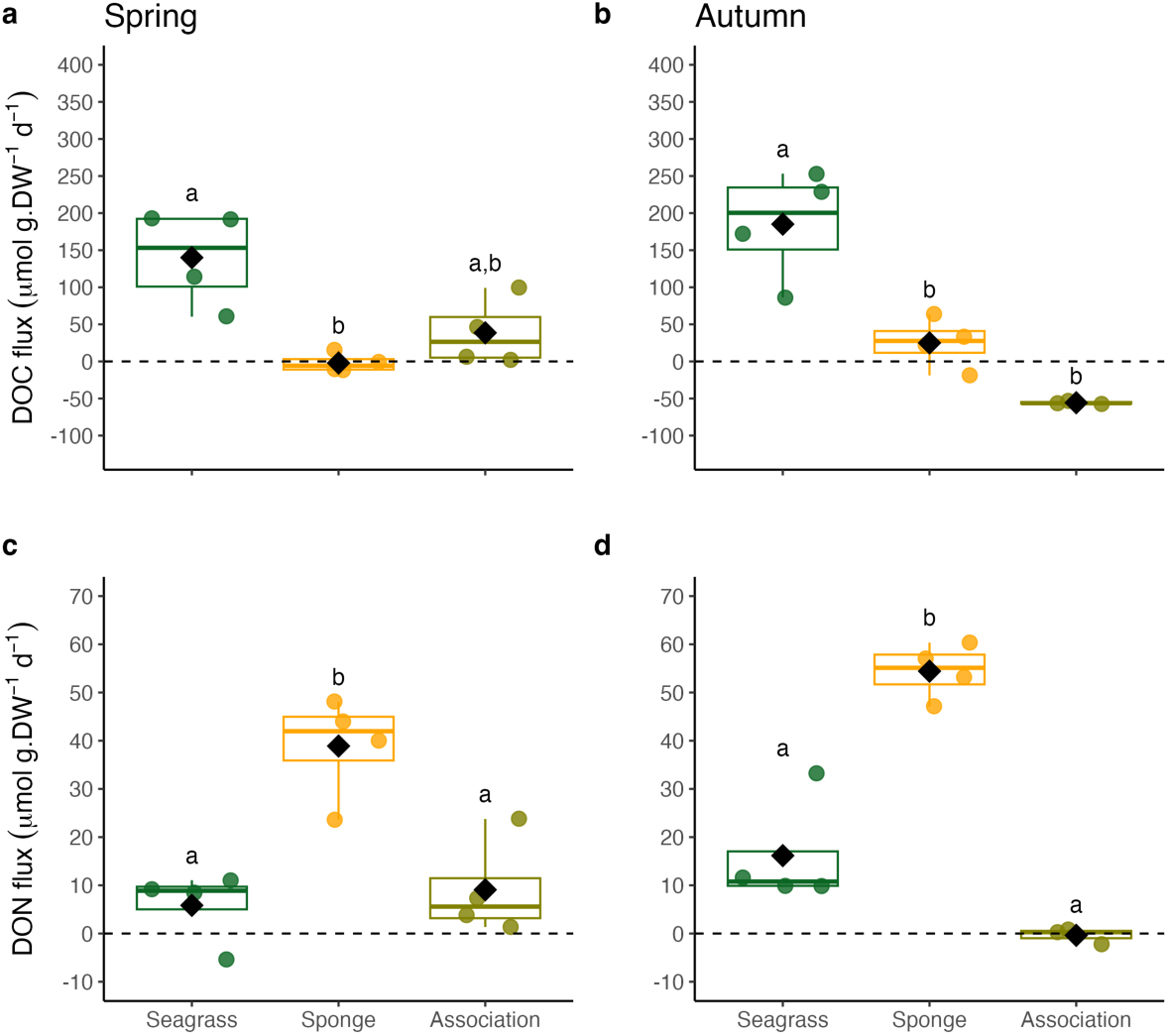
Daily fluxes of DOC (a, b) and DON (c, d) in incubations with the seagrass, sponge and the association in spring and autumn. Positive values indicate production, negative values indicate uptake. The center line denotes the median value (50^th^ percentile), the box contains the 25^th^ to 75^th^ percentiles. Whiskers mark the 5^th^ and 95^th^ percentiles. The horizontal dashed line marks the zero-flux threshold. Black squares indicate mean values; letters indicate significant differences between treatments, n=4.

### Primary production and respiration rates

Net community production (NCP) in spring was similar in the seagrass and the association (16.20 ± 2.48 and 19.24 ± 4.38 μmol g DW^-1^ h^-1^, respectively; mean ± SE, Fig. S2a, Tables S13, S14), followed by the sponge (1.79 ± 0.41 μmol g DW^-1^ h^-1^, Fig. S2a, Tables S13, S14). Community respiration (CR) in spring was highest (more negative) in the association (−4.63 ± 0.39 μmol g DW^-1^ h^-1^), followed by the sponge (3.92 ± 0.78 μmol g DW^-1^ h^- 1^), and the seagrass (1.79 ± 0.40 μmol g DW^-1^ h^-1^, Fig. S2b, Tables S13, S14). We observed a seasonal effect, where NCP in autumn was lower than in spring (Fig. S2c, Tables S13, S14), reaching 14.41 ± 3.35 μmol g DW^-1^ h^-1^ in the seagrass, 10.03 ± 1.35 μmol g DW^-1^ h^-1^ in the association and a net flux close to zero in the sponge. CR in autumn was higher (more negative) in autumn than in spring (Fig. S2d, Tables S13, S14), reaching −15.26 ± 2.22 μmol g DW^-1^ h^-1^ in the sponge, −7.77 ± 0.17 μmol g DW^-1^ h^-1^ in the association, and −5.32 ± 0.85 μmol g DW^-1^ h^-1^ in the seagrass.

### Microbial Community Structure

The 16s rRNA gene amplicon sequencing of the sponge-associated microbiome revealed a diverse microbial community (Fig. 4) differing between the water column and sponge community, but not between the sponges growing alone or in association with the seagrass (Fig. S3).

**Fig. 4.**
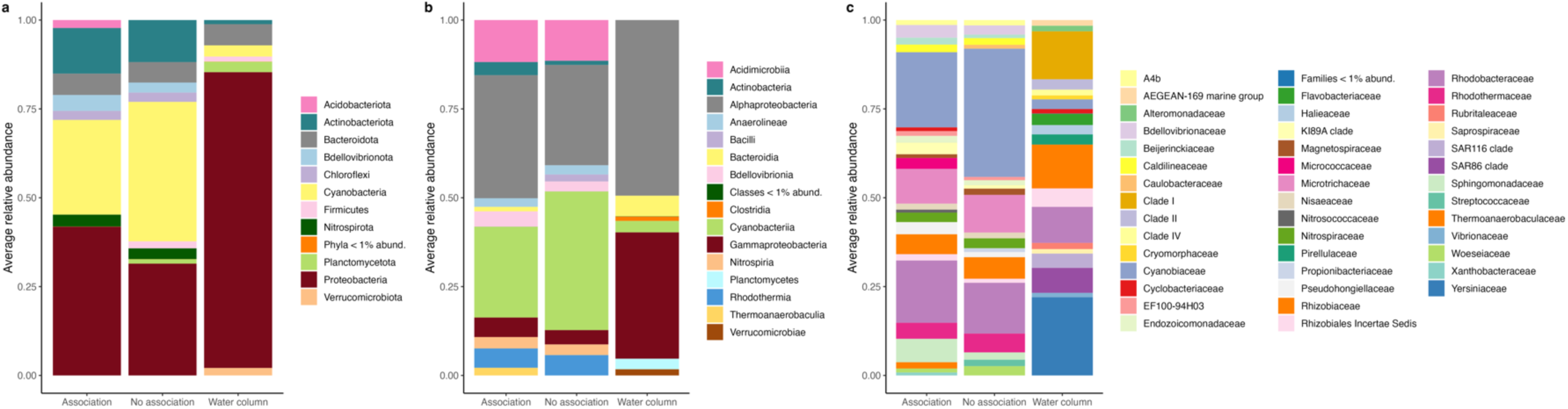
Average relative abundances of bacterial phyla (a), classes (b), and families (c) in *C. nucula* growing in association (n = 8), or growing alone (n = 4), and water column samples (n = 4).

The sponges were dominated by the phyla *Proteobacteria* (40.02 ± 4.26 %) and *Cyanobacteria* (32.17 ± 5.00 %) (Fig. 4). Among the predominant classes were *Alphaproteobacteria* (34.99 ± 3.98 %) and *Cyanobacteriia* (32.17 ± 5.00 %). Accordingly, the family of *Cyanobiaceae* accounted for 32.15 ± 4.99 % in sponge communities, followed by *Rhodobacteraceae* (20.52 ± 3.62 %) and *Microtrichaceae* (12.59 ± 1.28 %). The water column was dominated by the phylum of *Proteobacteria* (83.56 ± 0.96 %) with the classes *Alphaproteobacteria*(48.61 ± 4.83 %) and *Gammaproteobacteria* (34.95 ± 4.80 %). The most prevalent families were *Yersiniaceae* (20.75 ± 6.14 %) and Clade I (12.73 ± 2.99 %).

Taxonomic groups in the sponge microbiome with the largest effect detected in the differential abundance analysis (Fig. S4) were the *Synechococcus* spongiarum group within the family of *Cyanobiaceae* and *Silicimonas* within the family of *Rhodobacteraceae*, but also *Nitrospira* of the *Nitrospiraceae* family. Groups that had the highest effect on the differential abundance of the water column community were several species of the genus *Serratia* within the family of *Yersiniaceae*.

We found a significantly higher relative abundance of nitrifying families in the sponge communities compared to the water column (Table S15). No significant differences were found between the microbial communities of the sponge growing alone or in association. *Nitrospiraceae* were only found in sponge communities (Fig. 5) and accounted for 2.62 ± 0.76 % (mean ± SE). *Nitrosococcaceae* accounted for 0.42 ± 0.12 % of the sponge microbial communities and 0.09 ± 0.07 % in the water column (Fig. 5). Within the family of *Nitrosococcaceae* we found the genera Cm1-21 and AqS1, both belonging to AOB. Additionally, in a separate sequencing approach, we found the AOA taxon *Candidatus Nitrosopumilus* present in the sponge microbiome with no differences if the sponge was growing alone or in association with the seagrass.

**Fig. 5.**
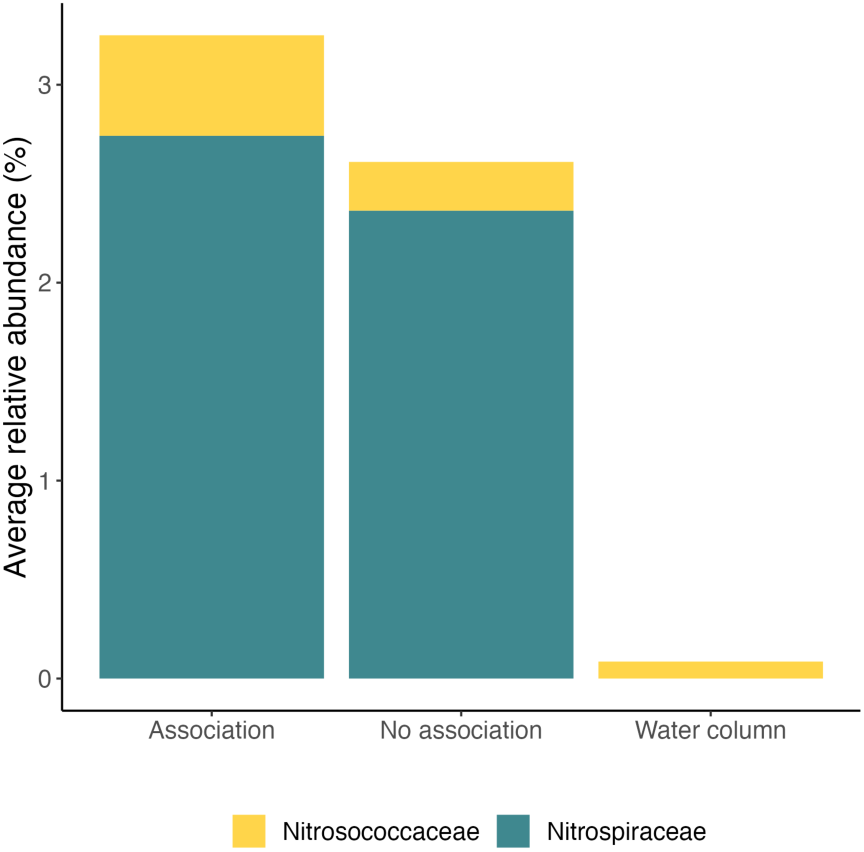
Average relative abundances of nitrifying families in the microbial community of *C. nucula* growing in association with *P. oceanica* (n=8), growing alone (n=4), and of the water column (n=4) in autumn.

## Discussion

This study of potential nitrification rates (PNR) and inorganic nutrient fluxes is the first to show that DIN provided by nitrification in the sponge *Chondrilla nucula* can be taken up by the seagrass *Posidonia oceanica*, supporting the N demand of the seagrass holobiont. The 16s rRNA gene amplicon sequencing of the sponge-associated microbiome revealed a diverse microbial community, including microorganisms involved in nitrification. Seagrasses in the Mediterranean Sea are threatened by climate change and anthropogenic impact (Boudouresque et al., 2009). This specific seagrass-sponge association, however, can thrive in an environment with high human pressure. Thus, investigating the biogeochemical and molecular mechanisms involved in its regulation can help to understand if this association can be beneficial for both partners under future environmental conditions.

### Nitrification contributes to N-cycling in the seagrass-holobiont

PNR of the *P. oceanica* - *C. nucula* association in dark incubations (21.51 ± 6.76 nmol g DW^-1^ h^-1^ in spring and 267.25 ± 33.01 nmol g DW^-1^ h^-1^ in autumn, Fig. 1) are within the range of those reported for other Mediterranean sponges (344 nmol g DW^-1^ h^-1^ in unstimulated incubations, Bayer et al., 2008; 180 - 780 nmol g DW^-1^ h^-1^, Jiménez & Ribes, 2007), and tropical sponges (30 - 2650 nmol g DW^-1^ h^-1^, Diaz & Ward, 1997). Reported PNR in surface sediments of seagrass meadows (0.15 - 1.0 μmol g^−1^ d^−1^, corresponding to 6.25 - 41.67 nmol g^−1^ h^−1^, Lin et al., 2021) or sandy estuaries (up to 40 nmol g DW^-1^ h^-^1 (Magalhães et al., 2005) are slightly lower. Since sponges are frequently found in many benthic environments such as seagrass meadows, the DIN excretion via nitrification can contribute significantly to nitrogen cycling (Diaz & Ward, 1997; Jiménez & Ribes, 2007). We found higher PNR in dark incubations, which is in line with the widely accepted explanation that both parts of the nitrification process (ammonium and nitrite oxidation) are light-inhibited (Guerrero & Jones, 1996; Horrigan et al., 1981).

We observed higher δ¹⁵N values in natural abundance samples of seagrass leaves and epiphytes when the plant was associated with the sponge in spring and of seagrass meristem tissue in autumn (Fig. S1). Nitrification is causing negative fractionation of NO_3_^-^ (depleted in δ¹⁵N) and positive fractionation of NH_4_^+^ (enriched in δ¹⁵N; Casciotti, 2016; Mariotti et al., 1981). NO_3_^-^ excreted from sponges and produced by nitrification, can therefore have lower δ¹⁵N values than NO_3_^-^ from the water column (Southwell, Popp, et al., 2008). At the same time, fractionation of NH_4_^+^ during uptake could increase the δ¹⁵N in the residual NH_4_^+^ pool in the sponge tissue (Hoch et al., 1994). Our higher δ¹⁵N values in plant and epiphyte tissue indicate that δ¹⁵N -enriched NH_4_^+^ excreted by the sponge was preferential taken up in the association. *P. oceanica* can assimilate N as NH_4_^+^ or NO_3_^-^ but usually shows a higher affinity for NH_4_^+^ (Touchette & Burkholder, 2000). Berlinghof et al. (2024) showed that *P. oceanica* prefers NH_4_^+^ uptake, while its epiphytes may preferentially use NO_3_^-^ as a strategy to avoid competition for N with the plant. The plant could therefore also compete for NH_4_^+^ with the sponge nitrifiers. We observed NH_4_^+^ production by the sponge only in spring, indicating that DIN excreted via nitrification might become more important in autumn. We measured high NO_3_^-^ production by the sponge (17.78 ± 3.49 μmol g DW^-1^ d^-1^ in spring and 23.59 ± 2.89 μmol g DW^-1^ d^-1^ in autumn), while the seagrass showed net fluxes close to zero or NO_3_^-^ uptake (Fig. 2). Net NO_3_^-^ fluxes in incubations with the seagrass-sponge association were also close to zero or showed NO_3_^-^ production, but lower compared to incubations with the sponge alone. This indicates that sponge-mediated nitrification produces NO_3_^-^ that is taken up by the seagrass holobiont. Whether NO_3_^-^ produced by sponge-associated nitrification benefits the seagrass or rather its associated epiphytes needs to be further investigated.

### Seasonal differences in PNR and nutrient fluxes

We observed PNR to be one order of magnitude higher in autumn than in spring (Fig. 1). Potential nitrification rates tend to be higher during the warmer seasons in salt marshes and estuary sediments. However, strong, site-specific variations are often reported (Caffrey et al., 2003; Dollhopf et al., 2005). While environmental conditions, such as temperature, light or water column O_2_ concentrations were similar across seasons (Table 1), ambient NH_4_^+^ concentrations were higher in spring (12.74 ± 3.09 μM in spring vs 2.76 ± 1.75 μM in autumn) and would therefore, in contrast to our findings, indicate higher PNR in spring.

*P. oceanica* can exhibit strong seasonal dynamics, depending on light and temperature, but also on local factors such as nutrient availability (Alcoverro et al., 1995). Metabolism studies show that the main growth phase occurs in spring while in autumn the seagrass is in a senescent phase (Berlinghof et al., 2022; Koopmans et al., 2020; Olivé et al., 2016). Accordingly, uptake rates of NH_4_^+^ and NO_3_^-^ by *P. oceanica* tend to be highest in spring and early summer (Lepoint et al., 2002; Nayar et al., 2010). At our study site, we observed seasonal morphological differences of the seagrass, indicating higher growth and biomass in spring and less in autumn, when we noticed high leaf loss and a shorter average leaf length. With *P. oceanica* being in its main growth phase in spring and early summer at our study location, there could have been increased competition for NH_4_^+^ by the plant, resulting in lower NH_4_^+^ availability for the nitrifying microbial community of the sponge and thus lower PNR.

### Potential effects at the holobiont and ecosystem level

We calculated the N demand of *P. oceanica* with the daily C budget based on the O_2_ fluxes and the C:N ratios of the holobiont (seagrass + epiphytes, Table 2). We further calculated the percentage of daily primary production of the seagrass holobiont that can be supported by sponge-mediated PNR. Based on these assumptions, sponge-mediated PNR can support 8.35 % of the holobiont primary production in spring and even 47.38 % in autumn. Since *P. oceanica* prefers the uptake of NH_4_^+^ over NO_3_^-^ (Touchette & Burkholder, 2000), while epiphytes potentially prefer NO_3_^-^ (Berlinghof et al., 2024), it appears as if mostly the seagrass epiphytes can benefit from sponge-mediated PNR.

**Table 2.** Nitrogen requirements of the *P. oceanica* holobiont (plant + epiphytes) in spring and autumn.

| Season | Daily C budget<br>($\mu\text{mol g DW}^{-1} \text{ d}^{-1}$) | CN ratio<br>holobiont | N demand<br>($\mu\text{mol g DW}^{-1} \text{ d}^{-1}$) | Daily PNR budget<br>($\mu\text{mol g DW}^{-1} \text{ d}^{-1}$) | % support |
| --- | --- | --- | --- | --- | --- |
| Spring | 57.37 | 23.3 | 2.462 | 0.21 | 8.35 |
| Autumn | 50.36 | 27.8 | 1.811 | 0.86 | 47.38 |

We observed DON production by *C. nucula* in both seasons (Fig. 3c, d). Ammonification of DON by seagrass-associated microbes produces NH_4_^+^ that can enhance the access of the seagrass to inorganic N as shown by Pfister et al., 2023. Thus, DON released by the sponge could further support the N demand of the seagrass holobiont. The sponge on the other side, can take up DOC release by the seagrass (Fig. 3a, b). However, the extent of these beneficial processes varies a lot throughout seasons and depends on environmental conditions, such as light or nutrient availability. Further investigations of the benefits for the sponge in this association are therefore needed.

### Microbial community

We found a diverse microbial community associated with *C. nucula* (Fig. 4) that was not affected by the association with *P. oceanica*. *Chondrilla nucula* harbors a distinct and stable bacterial community little affected by ambient seawater in the Mediterranean Sea (Thiel et al., 2007) or the Caribbean (Hill et al., 2006). Among the most prevalent and most distinct groups we found, was the *Synechococcus* spongiarum group (*Cyanobacteria*). These cyanobacterial symbionts are commonly found in marine sponges (Konstantinou et al., 2018; Usher, 2008), and are also reported for *C. nucula* (Usher et al., 2004). Another frequent group in the sponge communities were members of the family *Rhodobacteraceae* (*Alphaproteobacteria*); a family known to have several symbionts capable of fixing C via anoxygenic photosynthesis (Brinkmann et al., 2018) and also previously reported for *C. nucula* (Thiel et al., 2007). Families of the order Sphingomondales (*Alphaproteobacteria*) are known to be associated with marine sponges and have been linked to vitamin B12 synthesis (Thomas et al., 2010). Additionally, it has been demonstrated that the presence of *Sphingomonadaceae* can enhance degradation rates of artificial chemicals (Dai et al., 2022; Oh & Choi, 2019). As *C. nucula* has been shown to have a high capacity for bioaccumulation of pollutants (Ferrante et al., 2018), this could explain the high differential abundance of *Sphingomonas* in the microbial community of the sponge. Another dominant group was the family *Microtrichaceae*, which is potentially involved in nitrate supply as part of a nitrification-anammox system (Szitenberg et al., 2022).

Among the bacterial groups involved in nitrification (Fig. 5), we found the nitrite-oxidizing bacteria (NOB) *Nitrospira* within the family *Nitrospiraceae* (*Nitrospiria*), which is also reported for other species such as the cold-water sponge *Geodia baretti* (Hoffmann et al., 2009), but to our knowledge so far not for *C. nucula*. We also recovered the genera Cm1-21 and AqS1, both ammonia-oxidizing bacteria (AOB) within the family of *Nitrosococcaceae* (*Gammaproteobacteria*) (Hollingsworth et al., 2021; Semedo et al., 2021). We found the ammonia-oxidizing archaea (AOA) *Candidatus Nitrosopumilus*, and studies showed that they are stable associates of many sponge species (Bayer et al., 2008; Hoffmann et al., 2009; Holmes & Blanch, 2007).

Taken together, at the ecosystem level, we could show that the symbiosis between *P. oceanica*, *C. nucula*, and their microbiomes contributes significantly to nitrogen cycling. Since *C. nucula* shows a strong ability to compete for space (Bond & Harris, 1988; Milanese et al., 2003), it can quickly colonize new substrates. With increasing human pressure, that will open more space for the sponge to occupy (for example by boat anchoring in seagrass meadows, as we have seen at our study location), it is therefore essential to further investigate the dynamics of seagrass-sponge-associations and their implications at the ecosystem level as well as its potential as a nature-based solution for seagrass protection measures.

## Acknowledgments

We thank Paul Weber from the University of Bremen who assisted with the seagrass shoot density and sponge cover measurements. We thank L. Polaková and K. Umbria-Salinas of the SoWa Stable Isotope Facility (České Budějovice, CZ) for their assistance with nitrate isotope measurements. This research was supported by a Ph.D. fellowship co-funded by the Stazione Zoologica Anton Dohrn (SZN) and the University of Bremen (to J.B.) and a Ph.D. fellowship funded by the Open University – SZN Ph.D. Program (to L.M.M.). U.C. was partially supported by the Italian PRIN 2022 project ENGAGE (grant n. 20223R4FJK) and PRIN 2022 PNRR project BORIS (grant n. P2022R739J), funded by the European Union – Next Generation EU.

## Author contributions

J.B., L.M.M., C.W., and U.C. designed the study. J.B., L.M.M., and U.C. performed the experiments. T.B.M. performed the mass spectrometry analyses and J.B. and T.B.M. analyzed the results. J.B. and L.G. performed the molecular analyses and J.B. and H.G-V. analyzed the results. J.B. and U.C. wrote the manuscript with contributions from all co-authors.

