## Supplementary material for "Nitrification in a Seagrass-Sponge Association"

### Supplementary figures

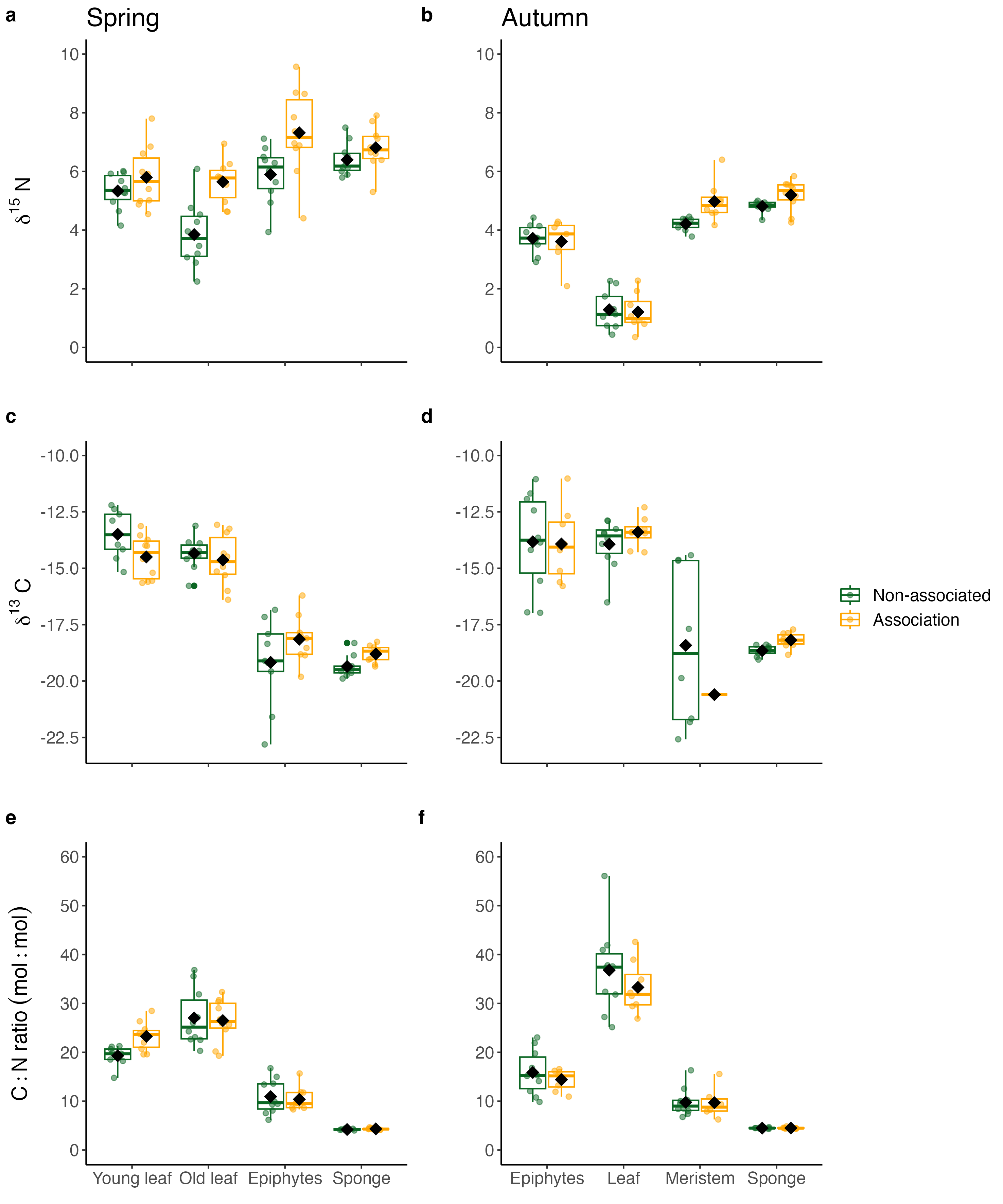

**Fig. S1.** Stable isotope composition and C/N ratios across communities in associated and non-associated states in spring (a, c, e) and autumn (b, d, f). (a, b) δ¹⁵N (‰), (c, d) δ¹^3^C (‰) and (e, f) C:N ratios for young and old Posidonia oceanica leaves, seagrass epiphytes and the sponge Chondrilla nucula. The center line denotes the median value (50^th^ percentile), the box contains the 25^th^ to 75^th^ percentiles. Whiskers mark the 5^th^ and 95^th^ percentiles. Black squares indicate mean values.

Spring

Autumn

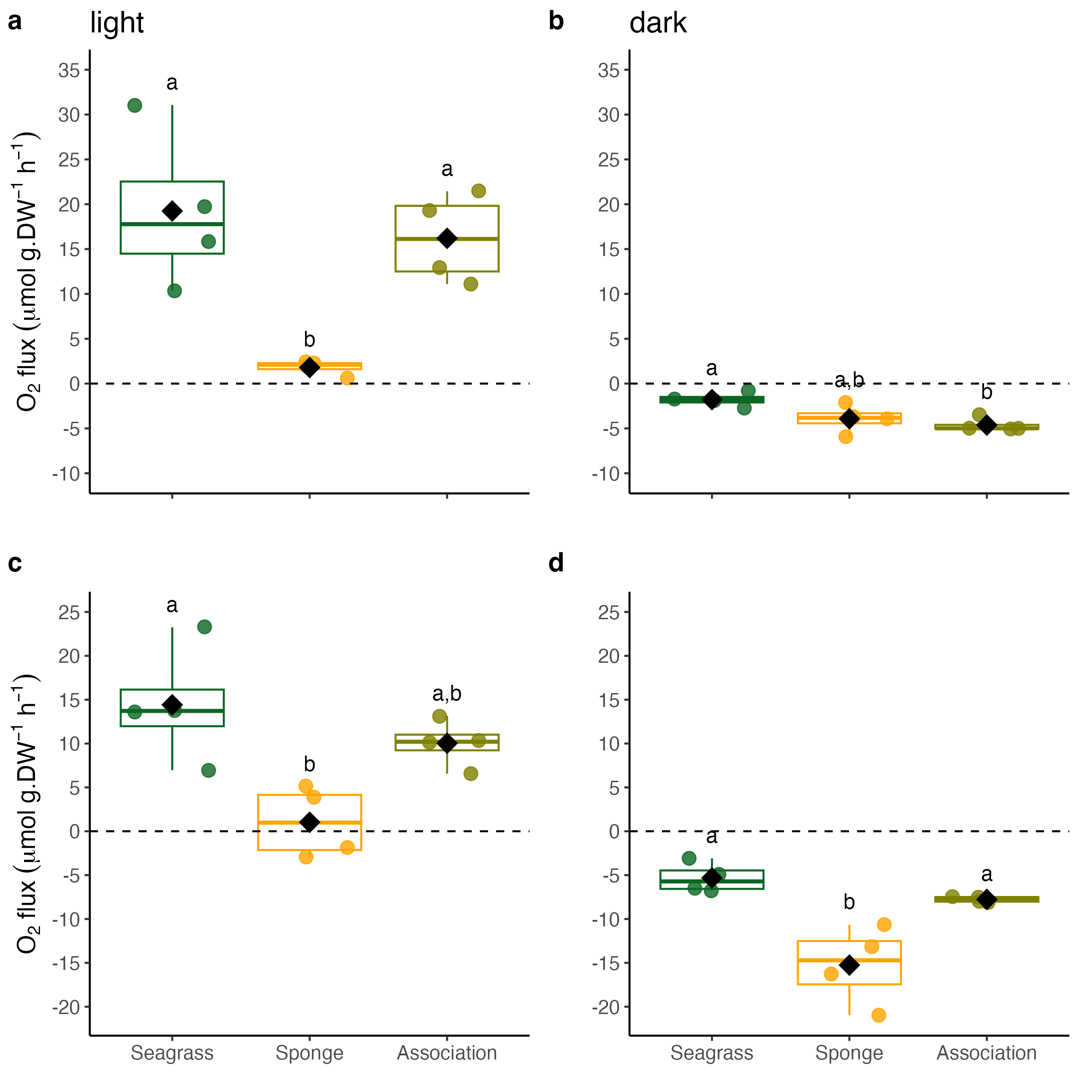

**Fig. S2.** O_2_ fluxes of the seagrass, sponge and the association. (a, c) Net community production (NCP) and (c, d) community respiration (CR) in incubations in spring (a, b) and autumn (c, d). Positive values indicate O_2_ production, negative values indicate respiration. The center line denotes the median value (50^th^ percentile), the box contains the 25^th^ to 75^th^ percentiles. Whiskers mark the 5^th^ and 95^th^ percentiles. The horizontal dashed line marks the zero-flux threshold. Black squares indicate mean values; letters indicate significant differences between treatments, n=4.

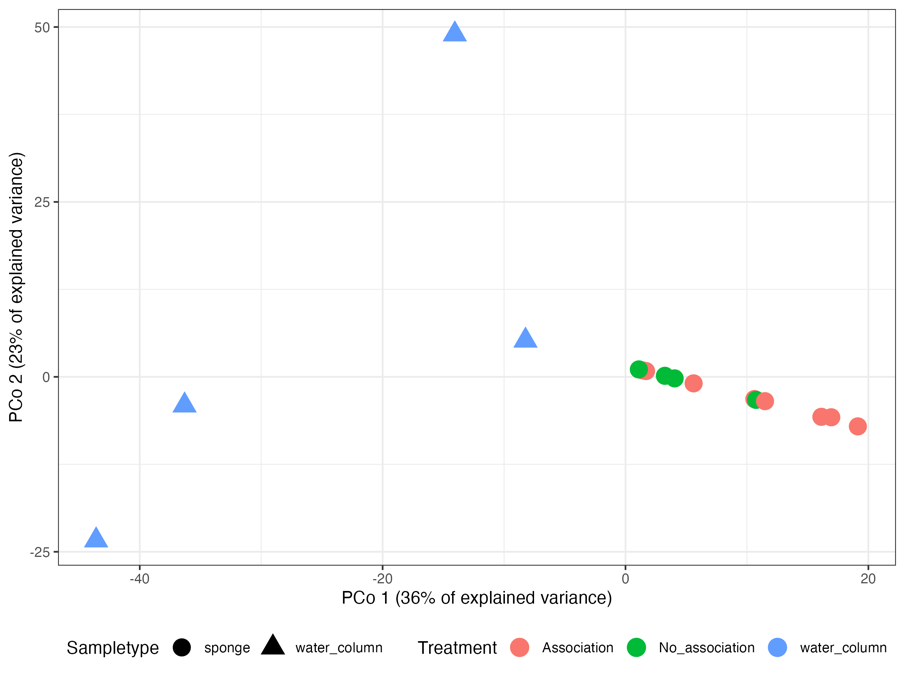

**Fig. S3.** Principal coordinates analysis of the bacterial community from *Chondrilla nucula* growing alone or in association and the water column community.

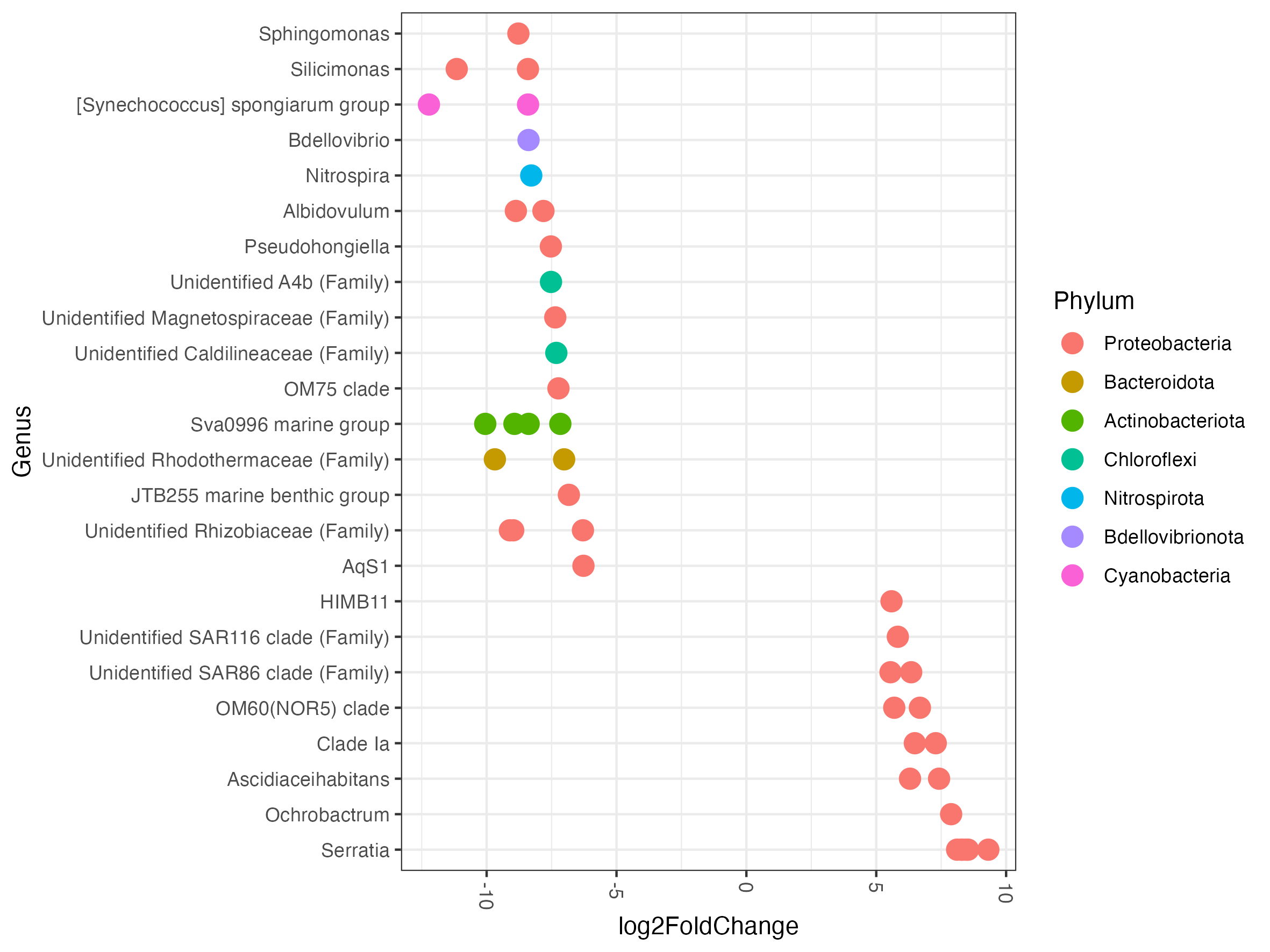

**Fig. S5.4.** Differential abundances in sponge and water column samples as log2FoldChange. Positive values mean differential abundance is higher in the water column and negative values higher in the sponges.

### Supplementary tables

**Table S1.** Permutation-based analysis of variance (PERMANOVA) for PNR (nmol N g DW⁻¹ h⁻¹) on different *community* types, *incubation* types, *seasons*, and their interactions. The table provides the degrees of freedom (Df), sum of squares (SS), proportion of variance explained (R²), pseudo-F statistics, and associated p-values (P(>F)) for each source of variation. Bold p-values (p < 0.05) indicate which factors contribute to differences in the measured variables.

| **Source of variation** | **Df** | **SS** | **R^2^** | **Pseudo-F** | **P (>F)** |
| --- | --- | --- | --- | --- | --- |
| Community | 2 | 79707 | 0.29 | 61.26 | **0.001** |
| Season | 1 | 58900 | 0.21 | 90.54 | **0.001** |
| Incubation | 1 | 16269 | 0.06 | 25.01 | **0.001** |
| Community: Season | 2 | 63357 | 0.23 | 48.70 | **0.001** |
| Community: Incubation | 2 | 16468 | 0.06 | 12.66 | **0.002** |
| Season: Incubation | 1 | 9010 | 0.03 | 13.85 | **0.001** |
| Community: Season: Incubation | 2 | 12072 | 0.04 | 9.28 | **0.002** |
| Residual | 31 | 20166 | 0.07 |  |  |
| Total | 42 | 275949 | 1.00 |  |  |

**Table S2.** Adjusted p-values from multilevel pairwise comparisons of PNR between *community* and *season*. The comparisons are performed using Tukey’s honest significant difference (HSD) test. Bold p-values indicate combinations that differ significantly (p < 0.05).

| **PNR** | Seagrass (spring) | Seagrass  (autumn) | Sponge (spring) | Sponge  (autumn) | Association (spring) | Association  (autumn) |
| --- | --- | --- | --- | --- | --- | --- |
| Seagrass (spring) | x |  |  |  |  |  |
| Seagrass (autumn) | 1.000 | x |  |  |  |  |
| Sponge (spring) | 0.988 | 0.999 | x |  |  |  |
| Sponge (autumn) | 0.637 | 0.724 | 0.857 | x |  |  |
| Association (spring) | 0.992 | 0.995 | 1.000 | 0.914 | x |  |
| Association (autumn) | **<0.001** | **<0.001** | **<0.001** | **<0.001** | **<0.001** | x |

**Table S3.** Adjusted p-values from multilevel pairwise comparisons of PNR between *community* and *incubation* type. The comparisons are performed using Tukey’s honest significant difference (HSD) test. Bold p-values indicate combinations that differ significantly (p < 0.05).

| **PNR** | Seagrass (light) | Seagrass  (dark) | Sponge (light) | Sponge  (dark) | Association (light) | Association  (dark) |
| --- | --- | --- | --- | --- | --- | --- |
| Seagrass (light) | x |  |  |  |  |  |
| Seagrass (dark) | 1.000 | x |  |  |  |  |
| Sponge (light) | 0.998 | 0.997 | x |  |  |  |
| Sponge (dark) | 0.971 | 0.956 | 0.999 | x |  |  |
| Association (light) | 0.688 | 0.595 | 0.863 | 0.967 | x |  |
| Association (dark) | **0.006** | **0.002** | **0.008** | **0.016** | 0.109 | x |

**Table S4.** Permutation-based analysis of variance (PERMANOVA) of δ¹⁵N (‰), δ¹^3^C (‰), and C:N ratio (mol:mol) on different *tissue* types, *association* types, *seasons*, and their interaction. The table provides the degrees of freedom (Df), sum of squares (SS), proportion of variance explained (R²), pseudo-F statistics, and associated p-values (P(>F)) for each source of variation. Bold p-values (p < 0.05) indicate which factors contribute to differences in the measured variables.

| **Variable** | **Source of variation** | **Df** | **SS** | **R^2^** | **Pseudo-F** | **P (>F)** |
| --- | --- | --- | --- | --- | --- | --- |
| **δ¹⁵N** | Tissue | 5 | 276.89 | 0.56 | 88.75 | **0.001** |
|  | Association | 1 | 18.50 | 0.04 | 29.65 | **0.001** |
|  | Season | 1 | 93.24 | 0.19 | 149.43 | **0.001** |
|  | Tissue: Association | 5 | 9.60 | 0.02 | 3.08 | **0.018** |
|  | Tissue: Season | 1 | 7.45 | 0.02 | 11.95 | **0.001** |
|  | Association: Season | 1 | 2.69 | 0.01 | 4.32 | **0.031** |
|  | Tissue: Association: Season | 1 | 2.63 | 0.01 | 4.22 | **0.045** |
|  | Residual | 136 | 82.99 | 0.17 |  |  |
|  | Total | 151 | 494.01 | 1.00 |  |  |
| **δ¹^3^C** | Tissue | 5 | 559.73 | 0.55 | 58.59 | **0.001** |
|  | Association | 1 | 0.02 | 0.00 | 0.01 | 0.925 |
|  | Season | 1 | 131.74 | 0.13 | 68.95 | **0.001** |
|  | Tissue: Association | 5 | 14.15 | 0.01 | 1.48 | 0.198 |
|  | Tissue: Season | 1 | 79.56 | 0.08 | 41.64 | **0.001** |
|  | Association: Season | 1 | 1.61 | 0.00 | 0.84 | 0.331 |
|  | Tissue: Association: Season | 1 | 1.22 | 0.00 | 0.64 | 0.420 |
|  | Residual | 124 | 236.93 | 0.23 |  |  |
|  | Total | 139 | 1024.97 | 1.00 |  |  |
| **C:N** | Tissue | 5 | 15645.9 | 0.87 | 211.15 | **0.001** |
|  | Association | 1 | 2.5 | 0.00 | 0.17 | 0.674 |
|  | Season | 1 | 104.1 | 0.01 | 7.02 | **0.010** |
|  | Tissue: Association | 5 | 139.9 | 0.01 | 1.89 | 0.115 |
|  | Tissue: Season | 1 | 86.3 | 0.01 | 5.82 | **0.017** |
|  | Association: Season | 1 | 1.2 | 0.00 | 0.08 | 0.773 |
|  | Tissue: Association: Season | 1 | 0.8 | 0.00 | 0.05 | 0.812 |
|  | Residual | 132 | 1956.2 | 0.11 |  |  |
|  | Total | 147 | 17936.8 | 1.00 |  |  |

**Table S5.** Adjusted p-values from multilevel pairwise comparisons of δ¹⁵N (‰) between *tissue* types and *association* types in spring. The comparisons are performed using Tukey’s honest significant difference (HSD) test. Bold p-values indicate combinations that differ significantly (p < 0.05).

| **δ¹⁵N** | Young leaves  (associated) | Young leaves (not associated) | Old leaves (associated) | Old leaves (not associated) | Epiphytes (associated) | Epiphytes (not associated) | Sponge (associated) | Sponge (not associated) |
| --- | --- | --- | --- | --- | --- | --- | --- | --- |
| Young leaves  (associated) | x |  |  |  |  |  |  |  |
| Young leaves (not associated) | 0.955 | x |  |  |  |  |  |  |
| Old leaves (associated) | 1.000 | 0.996 | x |  |  |  |  |  |
| Old leaves (not associated) | **<0.001** | **0.017** | **0.002** | x |  |  |  |  |
| Epiphytes (associated) | **0.014** | **<0.001** | **0.004** | **<0.001** | x |  |  |  |
| Epiphytes (not associated) | 1.000 | 0.887 | 0.999 | **<0.001** | **0.026** | x |  |  |
| Sponge (associated) | 0.267 | 0.018 | 0.126 | **<0.001** | 0.930 | 0.389 | x |  |
| Sponge (not associated) | 0.849 | 0.207 | 0.638 | **<0.001** | 0.384 | 0.932 | 0.977 | x |

**Table S6.** Adjusted p-values from multilevel pairwise comparisons of δ¹⁵N (‰) between *tissue* types and *association* types in autumn. The comparisons are performed using Tukey’s honest significant difference (HSD) test. Bold p-values indicate combinations that differ significantly (p < 0.05).

| **δ¹⁵N** | Leaves  (associated) | Leaves (not associated) | Epiphytes (associated) | Epiphytes (not associated) | Meristem (associated) | Meristem (not associated) | Sponge (associated) | Sponge (not associated) |
| --- | --- | --- | --- | --- | --- | --- | --- | --- |
| Leaves  (associated) | x |  |  |  |  |  |  |  |
| Leaves (not associated) | 1.000 | x |  |  |  |  |  |  |
| Epiphytes (associated) | **<0.001** | **<0.001** | x |  |  |  |  |  |
| Epiphytes (not associated) | **<0.001** | **<0.001** | 1.000 | x |  |  |  |  |
| Meristem (associated) | **<0.001** | **<0.001** | **<0.001** | **<0.001** | x |  |  |  |
| Meristem (not associated) | **<0.001** | **<0.001** | 0.369 | 0.517 | 0.092 | x |  |  |
| Sponge (associated) | **<0.001** | **<0.001** | **<0.001** | **<0.001** | 0.990 | **0.005** | x |  |
| Sponge (not associated) | **<0.001** | **<0.001** | **0.002** | **0.002** | 0.999 | 0.333 | 0.808 | x |

**Table S7.** Adjusted p-values from multilevel pairwise comparisons of δ¹^3^C (‰) between *tissue* types and *association* types in spring. The comparisons are performed using Tukey’s honest significant difference (HSD) test. Bold p-values indicate combinations that differ significantly (p < 0.05).

| **δ¹^3^C** | Young leaves  (associated) | Young leaves (not associated) | Old leaves (associated) | Old leaves (not associated) | Epiphytes (associated) | Epiphytes (not associated) | Sponge (associated) | Sponge (not associated) |
| --- | --- | --- | --- | --- | --- | --- | --- | --- |
| Young leaves  (associated) | x |  |  |  |  |  |  |  |
| Young leaves (not associated) | 0.452 | x |  |  |  |  |  |  |
| Old leaves (associated) | 1.000 | 0.292 | x |  |  |  |  |  |
| Old leaves (not associated) | 1.000 | 0.664 | 0.999 | x |  |  |  |  |
| Epiphytes (associated) | **<0.001** | **<0.001** | **<0.001** | **<0.001** | x |  |  |  |
| Epiphytes (not associated) | **<0.001** | **<0.001** | **<0.001** | **<0.001** | 0.461 | x |  |  |
| Sponge (associated) | **<0.001** | **<0.001** | **<0.001** | **<0.001** | 0.894 | 0.995 | x |  |
| Sponge (not associated) | **<0.001** | **<0.001** | **<0.001** | **<0.001** | 0.251 | 1.000 | 0.952 | x |

**Table S8.** Adjusted p-values from multilevel pairwise comparisons of δ¹^3^C (‰) between *tissue* types and *association* types in autumn. The comparisons are performed using Tukey’s honest significant difference (HSD) test. Bold p-values indicate combinations that differ significantly (p < 0.05).

| **δ¹^3^C** | Leaves  (associated) | Leaves (not associated) | Epiphytes (associated) | Epiphytes (not associated) | Meristem (associated) | Meristem (not associated) | Sponge (associated) | Sponge (not associated) |
| --- | --- | --- | --- | --- | --- | --- | --- | --- |
| Leaves  (associated) | x |  |  |  |  |  |  |  |
| Leaves (not associated) | 0.997 | x |  |  |  |  |  |  |
| Epiphytes (associated) | 0.998 | 1.000 | x |  |  |  |  |  |
| Epiphytes (not associated) | 0.999 | 1.000 | 1.000 | x |  |  |  |  |
| Meristem (associated) | **0.004** | **0.008** | **0.009** | **0.007** | x |  |  |  |
| Meristem (not associated) | **<0.001** | **<0.001** | **<0.001** | **<0.001** | 0.921 | x |  |  |
| Sponge (associated) | **<0.001** | **<0.001** | **<0.001** | **<0.001** | 0.867 | 1.000 | x |  |
| Sponge (not associated) | **<0.001** | **<0.001** | **<0.001** | **<0.001** | 0.953 | 1.000 | 0.998 | x |

**Table S9.** Permutation-based analysis of variance (PERMANOVA) of daily NH_4_^+^, NO_3_^-^ and NO_2_^-^ fluxes (μmol g DW^-1^ d^-1^) on different *community* types, *seasons*, and their interaction. The table provides the degrees of freedom (Df), sum of squares (SS), proportion of variance explained (R²), pseudo-F statistics, and associated p-values (P(>F)) for each source of variation. Bold p-values (p < 0.05) indicate which factors contribute to differences in the measured variables.

| **Variable** | **Source of variation** | **Df** | **SS** | **R^2^** | **Pseudo-F** | **P (>F)** |
| --- | --- | --- | --- | --- | --- | --- |
| **NH_4_^+^** | Community | 2 | 919 | 0.14 | 2.94 | 0.079 |
|  | Season | 1 | 2671 | 0.39 | 17.09 | **0.001** |
|  | Community: Season | 2 | 681 | 0.10 | 2.18 | **0.142** |
|  | Residual | 16 | 2501 | 0.37 |  |  |
|  | Total | 21 | 6771 | 1.00 |  |  |
| **NO_3_^-^** | Community | 2 | 2848 | 0.76 | 54.54 | **0.001** |
|  | Season | 1 | 5.4 | 0.00 | 0.21 | 0.651 |
|  | Community: Season | 2 | 400 | 0.11 | 7.66 | **0.003** |
|  | Residual | 18 | 470 | 0.13 |  |  |
|  | Total | 23 | 3724 | 1.00 |  |  |
| **NO_2_^-^** | Community | 2 | 0.39 | 0.27 | 9.98 | **0.002** |
|  | Season | 1 | 0.65 | 0.45 | 33.06 | **0.001** |
|  | Community: Season | 2 | 0.07 | 0.05 | 1.89 | 0.184 |
|  | Residual | 17 | 0.34 | 0.23 |  |  |
|  | Total | 22 | 1.49 | 1.00 |  |  |

**Table S10.** Adjusted p-values from multilevel pairwise comparisons of daily NH_4_^+^, NO_3_^-^ and NO_2_^-^ fluxes (μmol g DW^-1^ d^-1^) between *community* and *season*. The comparisons are performed using Tukey’s honest significant difference (HSD) test. Bold p-values indicate combinations that differ significantly (p < 0.05).

| **Variable** |  | Seagrass (spring) | Seagrass  (autumn) | Sponge (spring) | Sponge  (autumn) | Association (spring) | Association  (autumn) |
| --- | --- | --- | --- | --- | --- | --- | --- |
| **NH_4_^+^** | Seagrass (spring) | x |  |  |  |  |  |
|  | Seagrass (autumn) | 0.926 | x |  |  |  |  |
|  | Sponge (spring) | 0.077 | **0.012** | x |  |  |  |
|  | Sponge (autumn) | 0.941 | 1.000 | **0.021** | x |  |  |
|  | Association (spring) | 0.553 | 0.142 | 0.782 | 0.191 | x |  |
|  | Association (autumn) | 0.880 | 1.000 | **0.015** | 1.000 | 0.140 | x |
| **NO_3_^-^** | Seagrass (spring) | x |  |  |  |  |  |
|  | Seagrass (autumn) | 0.082 | x |  |  |  |  |
|  | Sponge (spring) | **<0.001** | **<0.001** | x |  |  |  |
|  | Sponge (autumn) | **<0.001** | **<0.001** | 0.604 | x |  |  |
|  | Association (spring) | 0.968 | **0.017** | **0.005** | **<0.001** | x |  |
|  | Association (autumn) | 0.089 | **<0.001** | 0.274 | **0.013** | 0.330 | x |
| **NO_2_^-^** | Seagrass (spring) | x |  |  |  |  |  |
|  | Seagrass (autumn) | 0.078 | x |  |  |  |  |
|  | Sponge (spring) | **0.047** | **<0.001** | x |  |  |  |
|  | Sponge (autumn) | 0.525 | 0.911 | **0.002** | x |  |  |
|  | Association (spring) | 0.946 | 0.340 | **0.008** | 0.934 | x |  |
|  | Association (autumn) | **0.045** | 1.000 | **<0.001** | 0.798 | 0.221 | x |

**Table S11.** Permutation-based analysis of variance (PERMANOVA) of daily DOC and DON fluxes (μmol g DW^-1^ d^-1^) on different *community* types, *seasons*, and their interaction. The table provides the degrees of freedom (Df), sum of squares (SS), proportion of variance explained (R²), pseudo-F statistics, and associated p-values (P(>F)) for each source of variation. Bold p-values (p < 0.05) indicate which factors contribute to differences in the measured variables.

| **Variable** | **Source of variation** | **Df** | **SS** | **R^2^** | **Pseudo-F** | **P (>F)** |
| --- | --- | --- | --- | --- | --- | --- |
| **DOC** | Community | 2 | 129813 | 0.69 | 28.25 | **0.001** |
|  | Season | 1 | 52 | 0.00 | 0.02 | 0.870 |
|  | Community: Season | 2 | 20703 | 0.11 | 4.51 | **0.039** |
|  | Residual | 17 | 39058 | 0.21 |  |  |
|  | Total | 22 | 189626 | 1.00 |  |  |
| **DON** | Community | 2 | 8131 | 0.85 | 14.33 | **0.001** |
|  | Season | 1 | 39 | 0.00 | 0.67 | 0.366 |
|  | Community: Season | 2 | 603 | 0.06 | 1.15 | **0.011** |
|  | Residual | 16 | 763 | 0.08 |  |  |
|  | Total | 21 | 9535 | 1.00 |  |  |

**Table S12.** Adjusted p-values from multilevel pairwise comparisons of daily DOC and DON fluxes (μmol g DW^-1^ d^-1^) between *community* and *season*. The comparisons are performed using Tukey’s honest significant difference (HSD) test. Bold p-values indicate combinations that differ significantly (p < 0.05).

| **Variable** |  | Seagrass (spring) | Seagrass  (autumn) | Sponge (spring) | Sponge  (autumn) | Association (spring) | Association  (autumn) |
| --- | --- | --- | --- | --- | --- | --- | --- |
| **DOC** | Seagrass (spring) | x |  |  |  |  |  |
|  | Seagrass (autumn) | 0.766 | x |  |  |  |  |
|  | Sponge (spring) | **0.007** | **<0.001** | x |  |  |  |
|  | Sponge (autumn) | **0.034** | **0.002** | 0.964 | x |  |  |
|  | Association (spring) | 0.074 | **0.005** | 0.831 | 0.998 | x |  |
|  | Association (autumn) | **<0.001** | **<0.001** | 0.690 | 0.286 | 0.157 | x |
| **DON** | Seagrass (spring) | x |  |  |  |  |  |
|  | Seagrass (autumn) | 1.000 | x |  |  |  |  |
|  | Sponge (spring) | **<0.001** | **<0.001** | x |  |  |  |
|  | Sponge (autumn) | **<0.001** | **<0.001** | 0.054 | x |  |  |
|  | Association (spring) | 1.000 | 1.000 | **<0.001** | **<0.001** | x |  |
|  | Association (autumn) | 0.500 | 0.411 | **<0.001** | **<0.001** | 0.481 | x |

**Table S13.** Permutation-based analysis of variance (PERMANOVA) of O_2_ fluxes (μmol g DW^-1^ d^-1^) on different *community* types, *incubation* types, *seasons*, and their interactions. The table provides the degrees of freedom (Df), sum of squares (SS), proportion of variance explained (R²), pseudo-F statistics, and associated p-values (P(>F)) for each source of variation. Bold p-values (p < 0.05) indicate which factors contribute to differences in the measured variables.

| **Source of variation** | **Df** | **SS** | **R^2^** | **Pseudo-F** | **P (>F)** |
| --- | --- | --- | --- | --- | --- |
| Community | 2 | 971.4 | 0.17 | 29.62 | **0.001** |
| Season | 1 | 295.7 | 0.05 | 18.03 | **0.001** |
| Incubation | 1 | 3426.4 | 0.61 | 208.98 | **0.001** |
| Community: Season | 2 | 7.5 | 0.00 | 0.23 | 0.791 |
| Community: Incubation | 2 | 211.1 | 0.04 | 6.44 | **0.005** |
| Season: Incubation | 1 | 13.0 | 0.00 | 0.79 | 0.387 |
| Community: Season: Incubation | 2 | 109.5 | 0.02 | 3.34 | **0.049** |
| Residual | 36 | 590.2 | 0.10 |  |  |
| Total | 47 | 5624.8 | 1.00 |  |  |

**Table S14.** Adjusted p-values from multilevel pairwise comparisons of O_2_ fluxes between *community* and *season*. The comparisons are performed using Tukey’s honest significant difference (HSD) test. Bold p-values indicate combinations that differ significantly (p < 0.05).

| **O_2_** | Seagrass (spring) | Seagrass  (autumn) | Sponge (spring) | Sponge  (autumn) | Association (spring) | Association  (autumn) |
| --- | --- | --- | --- | --- | --- | --- |
| Seagrass (spring) | x |  |  |  |  |  |
| Seagrass (autumn) | 0.962 | x |  |  |  |  |
| Sponge (spring) | 0.402 | 0.878 | x |  |  |  |
| Sponge (autumn) | **0.036** | 0.220 | 0.840 | x |  |  |
| Association (spring) | 0.992 | 1.000 | 0.758 | 0.137 | x |  |
| Association (autumn) | 0.670 | 0.984 | 0.998 | 0.590 | 0.940 | x |

**Table S15.** Permutation-based analysis of variance (PERMANOVA) of the nitrifying community abundance between sample type (sponge vs water column) and association type (associated vs non-associated). The table provides the degrees of freedom (Df), sum of squares (SS), proportion of variance explained (R²), pseudo-F statistics, and associated p-values (P(>F)) for each source of variation. Bold p-values (p < 0.05) indicate which factors contribute to differences in the measured variables.

| **Source of variation** | **Df** | **SS** | **R^2^** | **Pseudo-F** | **P(>F)** |
| --- | --- | --- | --- | --- | --- |
| Sample | 1 | 0.0013 | 0.11 | 3.5704 | **0.050** |
| Association | 1 | 0.0001 | 0.05 | 0.1492 | 0.729 |
| Residual | 29 | 0.0106 | 0.89 |  |  |
| Total | 31 | 0.0120 | 1.00 |  |  |
